# Size-dependent effect of site-specific polyethylene glycol modification of trastuzumab on the antibody functions

**DOI:** 10.64898/2026.09.23.753936

**Authors:** Yuya Tsutsui, Hiroaki Ohno, Hiroki Akiba

## Abstract

Polymer modification of antibodies has attracted attention for enhancing antibody functionality. Modification with polyethylene glycol (PEG) or biopolymers such as antibody-oligonucleotide conjugates has been investigated. Although direct effects at the modification site have been understood, distal effects of modifications on intrinsic antibody functions such as antigen-binding and Fc receptor-binding activities remain poorly characterized. In this study, we prepared site-specifically PEGylated trastuzumab to evaluate the effects of the modification site and the size of PEG chains on antibody functions. Reporter gene assays revealed a significant reduction in antibody-dependent cell cytotoxicity (ADCC) activity in a site- and size-dependent manner. Both the reduction in interaction with the antigen and FcγRIIIa were observed in a size-dependent manner. Interestingly, the effect of the modification site was only observed for ADCC activity, which suggests that the higher-order structure of the modified antibody plays a critical role in effector function. These results provide molecular design guidelines for the polymer modification of antibodies.

## Introduction

Chemical modification of antibodies is actively being studied, such as the development of advanced antibody-drug conjugates (ADCs)^1^. Polymer modification has been studied increasingly alongside small-molecule modifications. For instance, the introduction of polyethylene glycol (PEG) has been reported to improve the solubility and suppress the aggregation of ADCs loaded with hydrophobic drugs^2–6^. In these examples, PEG chains with molecular weights of several hundred have been commonly used. However, in recent years, studies utilizing polymers of kDa size have gained much attention for the efficient delivery of dozens of payloads. It has been reported that a branched PEG backbone reaching 200 kDa or a hydrophilic polypeptide exceeding 10 kDa counteracts drug hydrophobicity, thereby facilitating the achievement of high drug-to-antibody ratio^7,8^.

Furthermore, biopolymer-modified antibodies, including antibody-oligonucleotide conjugates (AOCs), are evolving rapidly^9^, with the molecular weight of the conjugated oligonucleotide reaching the kDa scale^10^. For instance, the introduction of a 40-kDa PEG into antisense oligonucleotides is known to improve serum stability and *in vivo* activity; thus, PEGylated AOCs have also been studied^11,12^. In addition to chemical modifications, protein fusions, such as bispecific antibodies and immunocytokines, have been widely developed as examples of site-specific modification^13–15^.

Such antibody modifications, however, may influence the biological functions of antibodies. Antibody-based therapeutics require the maintenance of antigen–antibody interactions. In addition, Fc receptor-mediated functions, including antibody-dependent cellular cytotoxicity (ADCC) and antibody-dependent cellular phagocytosis, play important roles in ADCs^16–18^. Thus, understanding the impact of polymer modification on antibody function is crucial for the rational design of antibody molecules.

It is well understood that polymer modification close to the antigen-binding surface can sterically hinder antigen binding, which has been successfully employed in prodrug strategies^19,20^. In addition, evaluations of the impact of random PEGylation via Lys residues within antibodies on antigen binding demonstrated that higher molecular weights of PEG and higher degrees of PEGylation lead to a decrease in both antigen-binding and Fc receptor-binding activities, accompanied by a reduction in effector functions^21^. Regarding the impact of site-specific PEGylation, previous evaluations have been performed on antigen-binding fragments (Fabs). It has been reported that site-specific PEGylation at the Cys residue in the hinge region of Fab’ did not affect its antigen-binding activity^22^. However, in full-size antibodies, the distal effects of site-specific modifications remain poorly understood.

We hypothesized that the modification site and molecular weight of the conjugates may influence the functions originating from molecular interactions, such as antigen binding and Fc receptor binding. In this study, we investigated these distal effects using trastuzumab, a monoclonal antibody against human epidermal growth factor receptor 2 (HER2), as our model. We systematically prepared site-specifically PEGylated trastuzumab variants to evaluate the effects of PEG modification site and molecular weight on antibody functions. By combining an ADCC reporter assays with antigen/FcγRIIIa binding measurements, we clarify the impact of polymer modification on FcγRIIIa- and antigen-binding activities.

## Results

### Preparation and Isolation of Site-Specific PEGylated Antibodies

Based on the THIOMAB technology^23^, genetically engineered Cys residues in heavy-chain H-A118C and light-chain L-V205C mutants (EU numbering) were site-specifically modified with PEGs of various sizes ranging from 6.9 nm (10 kDa) to 13 nm (40 kDa) as their hydrodynamic diameters (Figure 1)^24^. These mutants were selected based on their widespread use in site-specific modification^23,25,26^. Following reduction and re-oxidation treatments, the antibodies were reacted with maleimide-PEG reagents (Figure 2A). PEGylated antibodies were efficiently isolated by two-step cation-exchange chromatography (CIEX) (Figure 2B and Supporting Figure S1) and analyzed by SDS-PAGE (Figure 2C and Figure S2). Although reduced SDS-PAGE analysis revealed a residual unmodified light chain after the first CIEX, another round of CIEX effectively removed these unwanted species yielding sufficiently pure bis-PEGylated conjugates, the 40-kDa PEGylated L-V205C variant (Figure 2A and B). A similar observation was made for other modification campaigns (Supporting Figure S1 and S2).

**Figure 1.**
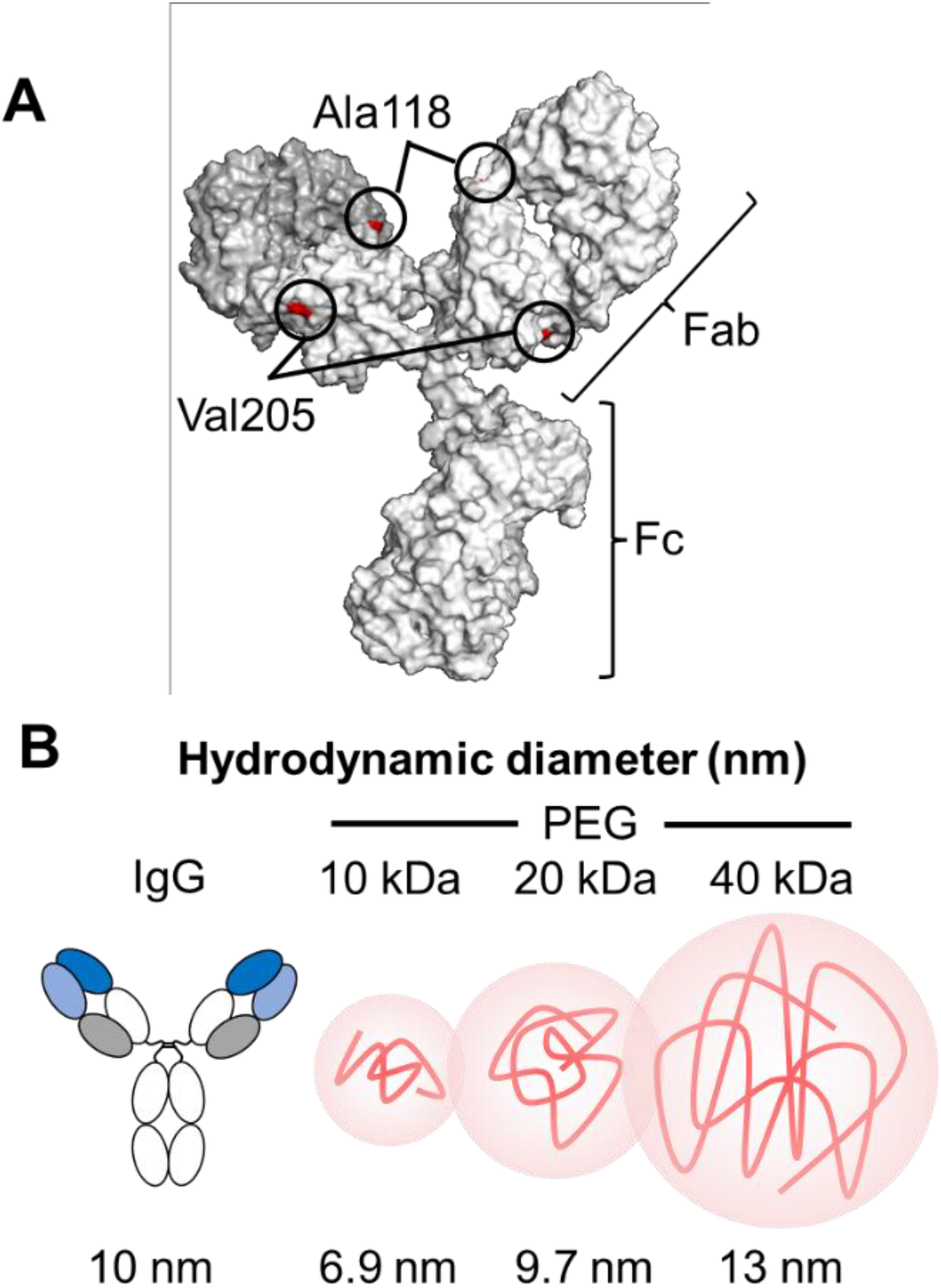
PEGylated-site and comparison of PEG-size depending on molecular weights. **(A)** Figure 1A shows schematic representation of the PEGylation sites in antibody. Site-specific PEGylation was conducted through engineered cysteine residues introduced at heavy chain A118C and light chain V205C. This model was created by molecular operating environment (MOE). **(B)** Figure 1B shows hydrodynamic diameter of PEG compared to IgG.

**Figure 2.**
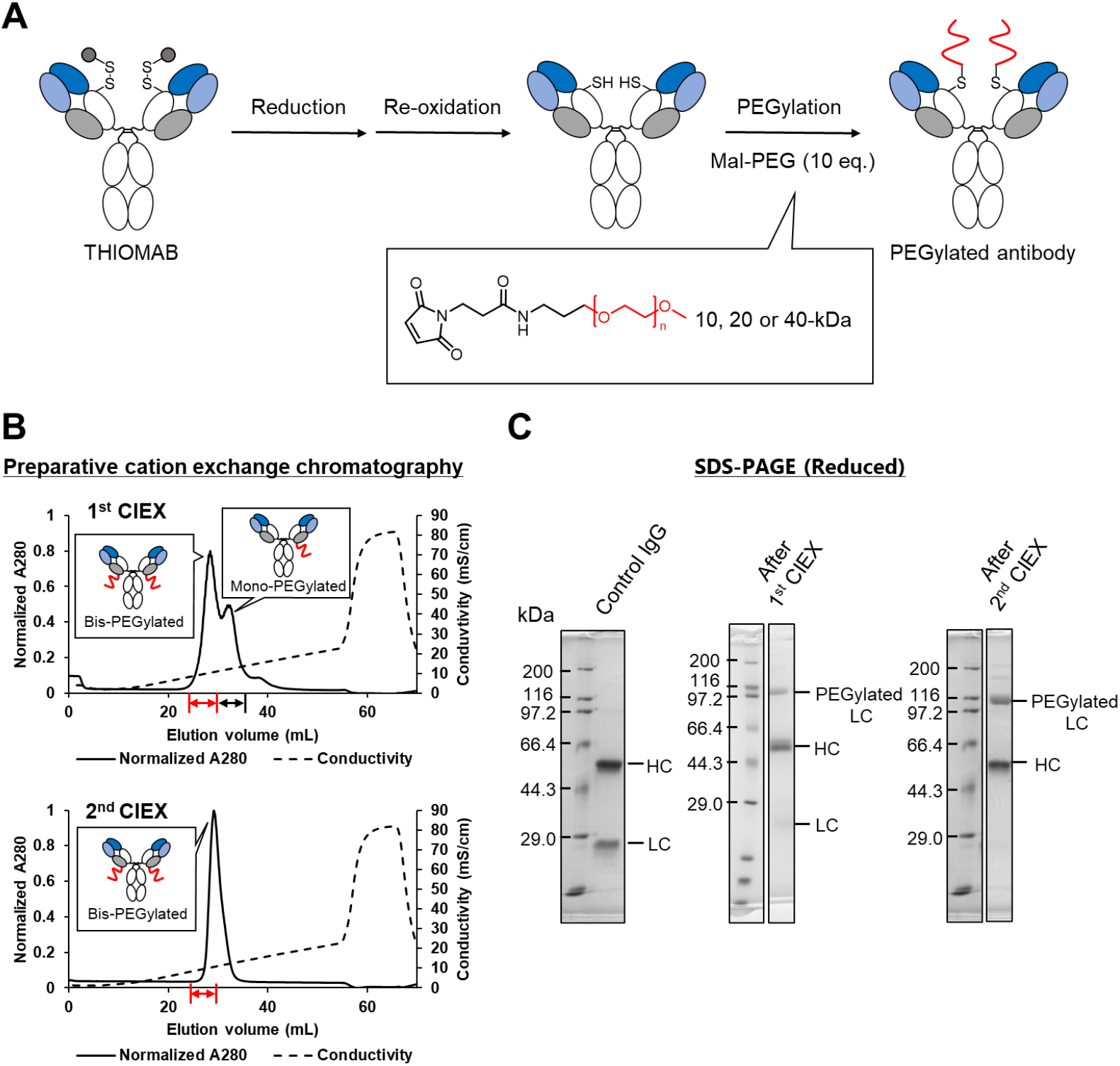
Synthesis and purification of PEGylated antibodies. **(A)** THIOMABs were reduced using dithiothreitol and reoxidized by air oxidation. Reoxidized THIOMABs were then reacted with maleimide-PEG (Mal-PEG) reagents. **(B)** Purification strategy for L-V205C conjugated with 40-kDa PEG using two-step cation exchange chromatography (CIEX). In the 1^st^ CIEX, fractions for bis-PEGylated antibodies (red) were collected, whereas those for mono-PEGylated antibodies (black) were discarded. The collected fractions were subjected to 2nd CIEX, and the red fractions were collected. Solid lines, absorbance at 280 nm (A280); dashed lines, electrical conductivity. **(C)** Samples after the 1^st^ and 2^nd^ CIEX were analyzed by reduced SDS-PAGE. A non-PEGylated antibody was used as a control (Control IgG).

To confirm that the site-specific conjugation of PEG chains did not induce global structural changes or misfolding, the thermal stability of the obtained PEGylated variants was evaluated using differential scanning fluorimetry. No significant differences were observed between the wild-type (WT) and any of the PEGylated variants; the transition temperatures *T*_m1_ (denaturation of the CH2 domains) and *T*_m2_ (denaturation of the Fab fragment) remained virtually unchanged at around 66 °C and 83 °C upon modification. This indicated that the effect of PEGylation on stability was negligible in the following experiments (Figure S3, Table S1).

### ADCC Activities

The effect of PEGylation on antibody-dependent cellular cytotoxicity (ADCC) activity was evaluated using a reporter gene assay system expressing FcγIIIa by co-culturing with the HER2-positive cell line SK-BR-3 (Figure 3). For the heavy chain variant H-A118C, we observed a PEG-size-dependent decrease in the maximum response (E_max_) of nuclear factor of activated T-cells (NFAT) transcription downstream of FcγIIIa (Table 1). Specifically, as the PEG molecular weight increased from 10- to 40-kDa, the E_max_ dropped markedly compared to that of the wild-type (WT) antibody. In contrast, the effect of the light chain variant L-V205C was less significant, with the 10-kDa PEGylated variant exhibiting an E_max_ value comparable to that of the WT, while only the larger 20-kDa and 40-kDa variants showed decreased in a size-dependent manner. For both mutants, the EC_50_ values increased slightly depending on the molecular weight of PEG, indicating a size-dependent reduction in signaling potency (Table 1). Taken together, these results demonstrate that both the molecular weight and modification site of PEG influence ADCC activity.

**Table 1.** Summary of parameters.

|  |  | ADCC reporter assay |  | Cell | Antigen binding in SPR |  |  | Retention time in FcγRIIIa |
| --- | --- | --- | --- | --- | --- | --- | --- | --- |
|  |  | EC<sub>50</sub> (nM)<sup>a</sup> | E<sub>max</sub> (nM) | binding | k<sub>on</sub> | k<sub>off</sub> (1/s) | K<sub>D</sub> | affinity chromatography |
|  |  |  |  | EC<sub>50</sub> (nM)<sup>a</sup> | (1/Ms) |  | (nM) | (min) |
| WT |  | 0.336 ± 0.035 | 1.02 ± 0.01 | 3.01 ± 0.32 | 3.10×10<sup>5</sup> | 5.07×10<sup>-4</sup> | 1.64 | 11.9 |
| H-A118C | 10-kDa | 1.27 ± 0.10 | 0.680 ± 0.065 | 7.68 ± 0.52 | 3.27×10<sup>5</sup> | 4.22×10<sup>-4</sup> | 1.19 | 9.39 |
|  | 20-kDa | 4.05 ± 0.48 | 0.299 ± 0.080 | 15.5 ± 0.9 | 3.50×10<sup>5</sup> | 5.07×10<sup>-4</sup> | 1.21 | 3.36 |
|  | 40-kDa | ≥25.7 | 0.115 ± 0.032 | 19.3 ± 0.8 | 3.30×10<sup>5</sup> | 3.89×10<sup>-4</sup> | 1.18 | 1.79 |
| L-V205C | 10-kDa | 0.750 ± 0.173 | 1.07 ± 0.13 | 8.79 ± 0.66 | 3.97×10<sup>5</sup> | 5.04×10<sup>-4</sup> | 1.27 | 8.57 |
|  | 20-kDa | 2.72 ± 0.48 | 0.763 ± 0.063 | 15.3 ± 1.1 | 3.77×10<sup>5</sup> | 5.10×10<sup>-4</sup> | 1.35 | 3.41 |
|  | 40-kDa | 7.78 ± 0.41 | 0.456 ± 0.027 | 26.3 ± 1.5 | 4.27×10<sup>5</sup> | 6.08×10<sup>-4</sup> | 1.42 | 2.78 |

<sup>a</sup> mean ± SD.

**Figure 3.**
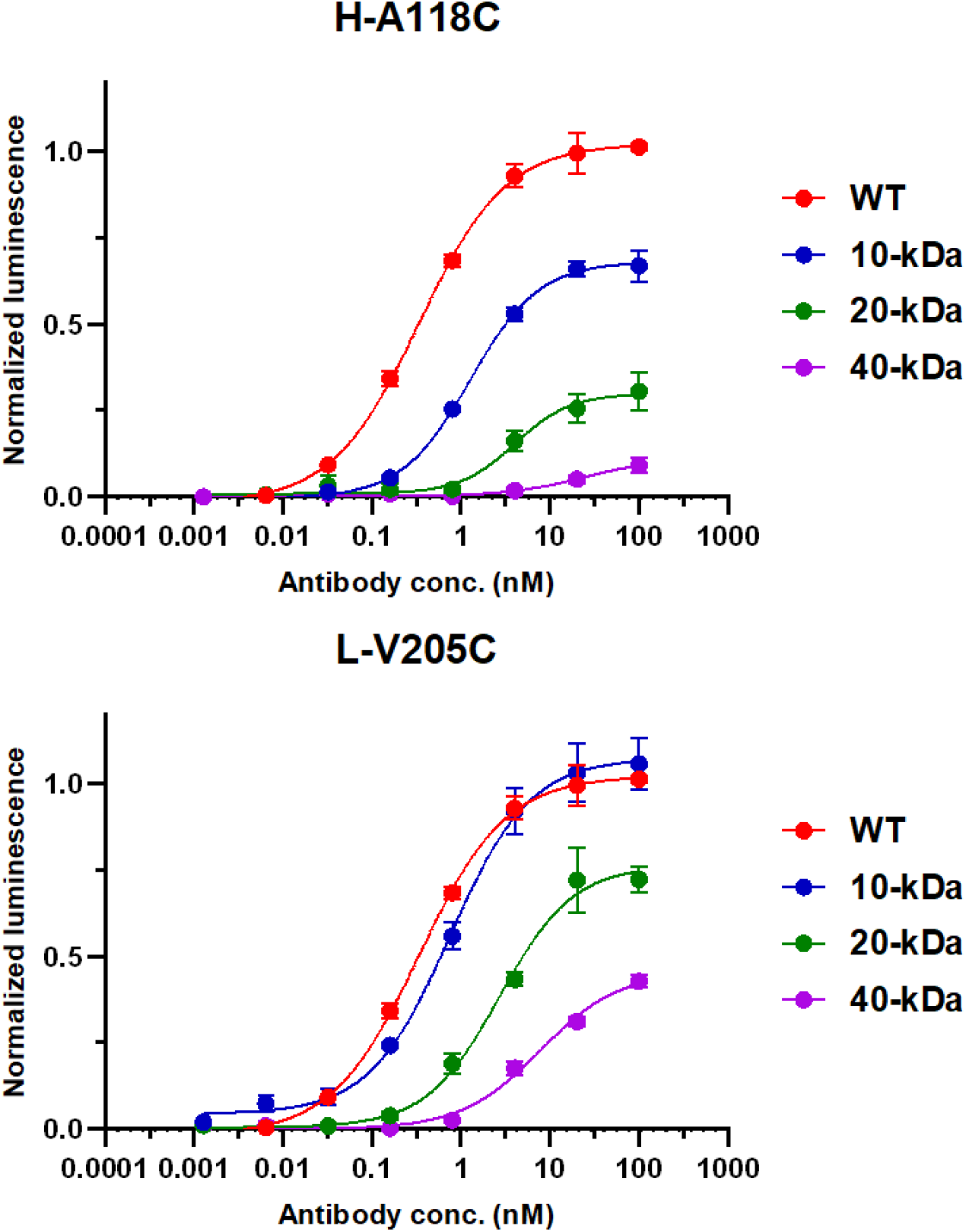
Antibody-dependent cellular cytotoxicity (ADCC) activity of PEGylated antibodies. The ADCC activity of the antibodies was evaluated using NFAT luciferase reporter assays based on FcγRIIIa-dependent signal transduction. ADCC activity was quantified by measuring the enzymatic activity of luciferase, which was expressed as a reporter gene. Data are shown as mean ± standard deviation from three to four experiments.

### FcγRIIIa-Binding Activities

The decrease in ADCC activity of the PEGylated variants was hypothesized to stem from a reduction in their FcγRIIIa-binding activity. The binding of WT and PEGylated antibodies using FcγRIIIa affinity chromatography (Figure 4). Both the H-A118C and L-V205C mutants exhibited a PEG size-dependent decrease in retention time. Even the smallest 10-kDa PEG possesses a hydrodynamic diameter of approximately 7 nm, making the PEG molecule comparable in hydrodynamic size to Fab fragment^24^. Thus, the introduced PEG would sterically hinder FcγRIIIa binding, leading to a size-dependent decrease in affinity, resulting in a size-dependent reduction in affinity and shorter retention time. Notably, the chromatograms of the WT and the 10- and 20-kDa PEGylated variants of both mutants displayed three distinct peaks. This profile can be attributed to differences in binding activity arising from variations in the N-linked glycan structure at Asn297^27^.

**Figure 4.**
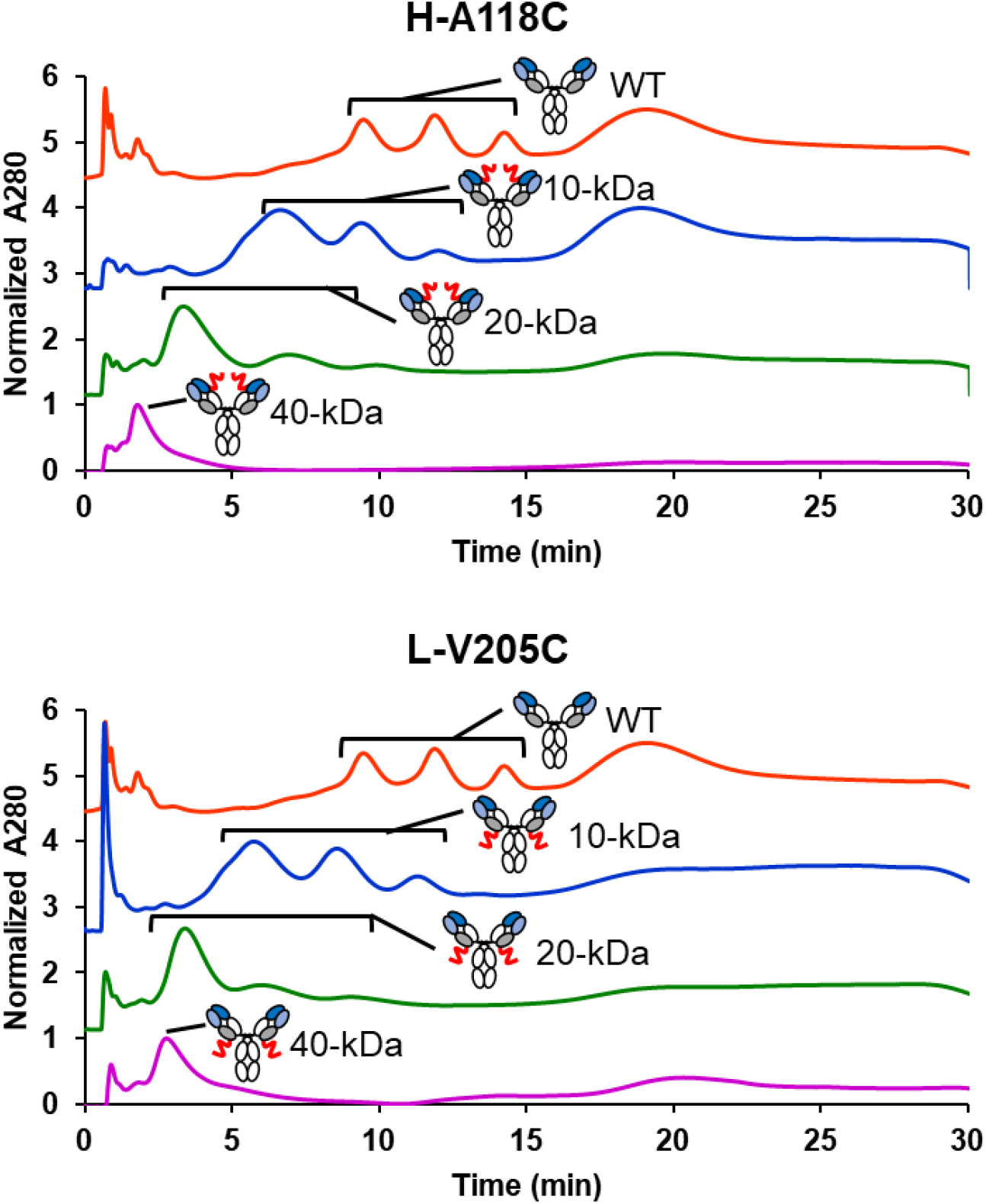
Normalized chromatograms of WT and PEGylated antibodies using an FcγRIIIa affinity column.

Intriguingly, while a molecular weight-dependent decrease in FcγRIIIa-binding activity was observed, no differences were detected between the two modification sites, indicating that evaluating FcγRIIIa-binding activity alone is insufficient to fully explain the varying degrees of ADCC activity reduction.

### Antigen-Binding Activities

In addition to FcγR binding, antigen binding is essential for ADCC. To assess the impact of site-specific PEGylation on target engagement, we evaluated the binding activities of the wild-type and PEGylated antibodies to HER2 on SK-BR-3 cell membranes using flow cytometry (Figure 5A). The EC_50_ values were calculated from the resulting sigmoidal curves (Table 1). The WT antibody exhibited an EC_50_ of 3.0 nM, which was in close agreement with previously reported values (approximately 1 nM)^28,29^. For PEGylated variants, the EC_50_ values increased in a PEG molecular weight-dependent manner, confirming that conjugation of larger PEG chains progressively impairs antigen-binding activity.

**Figure 5.**
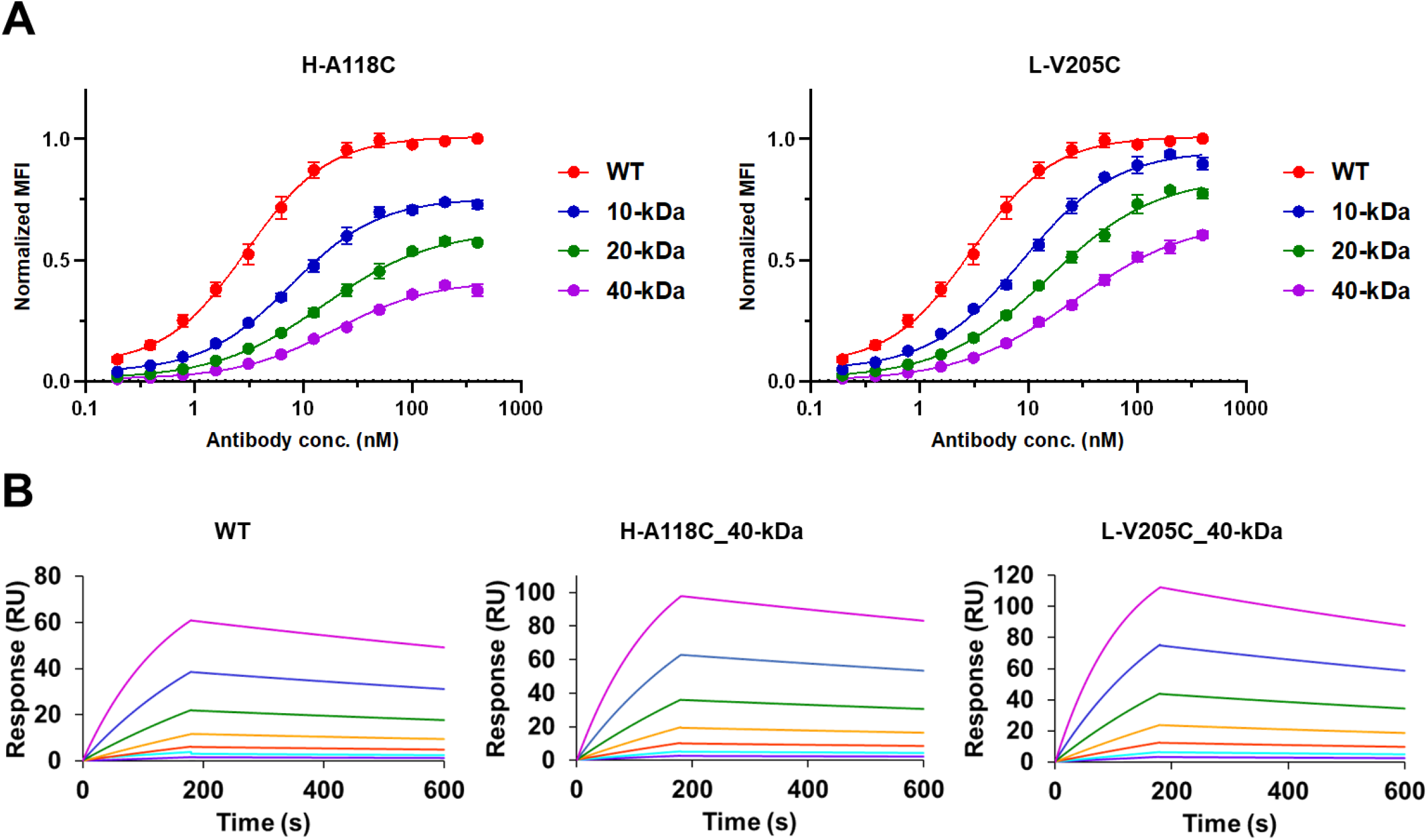
Antigen binding of PEGylated antibodies. **(A)** Cell binding of WT and PEGylated antibodies quantified by mean fluorescence intensity (MFI) using flow cytometry. Fitted curves were obtained from a four-parameter logistic model and are presented as mean ± SD from three experiments. **(B)** Surface plasmon resonance sensorgrams of immobilized WT and PEGylated antibodies binding to HER2.

Interestingly, antibody-captured surface plasmon resonance (SPR) measurements demonstrated that the affinity of each variable region was not affected by PEGylation (Figure 5B and Figure S4). Kinetic parameters (*k*_*on*_ and *k*_*off*_) for the WT were generally consistent with reported values (Table 1)^30,31^. The values for the PEGylated variants were comparable to those of WT, regardless of the PEG size or modification site. These SPR results clearly demonstrate that PEGylation of the antibody does not affect the monovalent affinity.

To understand the differences between the cell-surface binding and SPR analyses, bivalent binding of antibodies to cellular antigens should be considered: while the SPR evaluated monovalent binding, cell-surface target engagement involves bivalent interactions. Therefore, the size-dependent reduction in cell-binding affinity suggests that PEGylation affects the cooperative bivalent engagement of the two Fab fragments with adjacent cell-surface antigens.

## Discussion

In this study, we conducted a cross-validation of the size and position of site-specific PEG introduced into antibody Fabs. Our analyses revealed that the reduction in ADCC activity with increasing PEG size could not be explained independently by the reduction in either FcγRIIIa or antigen-binding activities. Effective ADCC activity requires a cooperative ternary interaction among the target antigen, antibody, and Fc receptor^32^. Beyond individual binary complex formation, PEGylation interferes with cooperative molecular recognition during the formation of the ternary antigen–antibody–Fc receptor complex. As a low-affinity Fc receptor, FcγRIIIa relies on ternary clustering to promote immunological synapse formation and effective signaling. In our cellular study, the decrease in Emax strongly suggested the inhibition of this cluster formation by PEGylation, which may be partly explained by the position-dependent differences. According to the structural model of the trastuzumab–HER2 complex^33^, the antibody light chain is oriented toward the membrane-proximal position, whereas the heavy chain points away from it. Thus, the introduction of PEG to the light chain may affect HER2-expressing cell binding but not FcγR-mediated cluster formation. In contrast, heavy-chain PEGylation (H-A118C) affects the density of receptor complexes in cell-cell interactions, explaining why Emax was more severely affected by PEGylation in H-A118C than in L-V205C.

In conclusion, this study successfully established a robust platform for the preparation and isolation of highly homogeneous, site-specifically PEGylated trastuzumab variants. Our findings demonstrated that site-specific PEGylation reduces both FcγRIIIa-binding and bivalent antigen-binding activities in a size-dependent manner, thereby decreasing ADCC activity. Specifically, PEGylation sterically interferes with the cooperative molecular recognition required to assemble of the antigen–antibody–Fc receptor complex, hindering cluster formation at the cell–cell interface. These insights elucidate the cooperative influence of modification sites and polymer molecular weights, providing vital design principles for the rational molecular design of antibody-based therapeutics.

### Experimental Procedures Cell culture

As an HER2-expressing cell line, SK-BR-3 cells were cultured in RPMI1640 containing 10% fetal bovine serum (FBS) and 1% penicillin/streptomycin. Jurkat-Lucia^TM^ NFAT-CD16 cells (Invivogen), which are NFAT-driven luciferase reporters, were cultured in IMDM containing 2 mM glutamine, 10% FBS, 1% penicillin/streptomycin, 100 µg/mL blasticidin, and 1 mg/mL zeocin.

### Expression and purification of wild-type trastuzumab and THIOMABs

Cysteine mutations were introduced into the antibody light or heavy chain by PCR-based site-directed mutagenesis using the KOD-Plus-Mutagenesis Kit (TOYOBO). Plasmids of wild-type trastuzumab and THIOMABs in the pcDNA3.4 vector were transfected into the Expi293F expression system (Thermo Fisher Scientific) according to the manufacturer’s standard protocol. The cells were removed by centrifugation at 6000 × g for 15 min, and the culture supernatant was filtered through a 0.20 μm filter. Antibodies were captured using an rProtein A Sepharose Fast Flow column (Cytiva, Tokyo, Japan). The column was washed with 10 mL of phosphate-buffered saline (PBS; pH 7.4) and eluted with 100 mM sodium citrate, pH 3.0 (3.0 mL). The eluent was dialyzed into PBS and purified by size-exclusion chromatography using a HiLoad 16/600 Superdex 200 pg column (Cytiva) in PBS.

### Expression, and purification of HER2-MBP

Maltose-binding protein-fused HER2 (HER2-MBP) C-terminally tagged with hexahistidine was expressed in the Expi293F expression system. After 7 days of culture, the supernatant was filtered through a 0.20 μm filter and then dialyzed into buffer A (20 mM Tris, 300 mM NaCl, pH 8.0). Ni^2+^ affinity chromatography was conducted using a HisTrap^TM^ HP column (Cytiva) with buffer A and buffer B (20 mM Tris, 300 mM NaCl, 500 mM imidazole, pH 8.0). HER2-MBP was eluted using a linear gradient from 0%B to 100%B for 20 min. The eluent was dialyzed into buffer C (20 mM Tris, 10 mM NaCl, pH 8.0). Anion exchange chromatography was conducted using a Capto^TM^ HiRes Q column with buffer C (20 mM Tris, 10 mM NaCl, pH 8.0) and buffer D (20 mM Tris, 1 M NaCl, pH 8.0). HER2-MBP was eluted using a linear gradient from 0%D to 50%D for 50 min. The fraction containing HER2-MBP was subjected to final purification using a HiLoad 16/600 Superdex 200 pg column (Cytiva) in buffer A.

### Bioconjugation of PEGylated antibody

THIOMAB (36 µM) was reduced using dithiothreitol (DTT, 3.6 mM) in PBS for 2 h at 37 °C. DTT was removed using PD MiniTrap^TM^ G-25 (Cytiva). The antibody solution was diluted to 0.25 µM and incubated for 3.5 h at 25 °C for re-oxidation. Maleimide-functionalized polyethylene glycol (Mal-PEG; SUNBRIGHT MA series, NOF CORPORATION) was dissolved in PBS to a concentration of 25 µM. The Mal-PEG solution was added to the antibody solution at a 10% (v/v) ratio, followed by the addition of EDTA to a final concentration of 1 mM. The reaction mixture was incubated for 2 h at 37 °C.

### Purification of PEGylated antibodies by cation exchange chromatography

PEGylated antibodies were purified by cation-exchange chromatography (CIEX) using a Capto^TM^ HiRes S 5/50 GL column (Cytiva) with buffer A (20 mM MES-NaOH, 10 mM NaCl, pH 5.0) and buffer B (20 mM MES-NaOH, 1 M NaCl, pH 5.0). PEGylated antibodies were eluted using a linear gradient from 0% B to 25% B for 50 min. The PEGylated antibodies were dialyzed and stored in PBS.

### Differential Scanning Fluorimetry

Differential scanning fluorimetry was conducted using the Protein Thermal Shift^TM^ Dye kit (Thermo Fisher Scientific), according to the manufacturer’s standard protocol. Antibody samples were prepared in PBS at a final concentration of 8 µM in a total volume of 12.5 µL. The antibody solution was mixed with 5 µL of Protein Thermal Shift buffer and 2.5 µL of diluted Protein Thermal Shift™ Dye (8 ×). Measurements were performed using a real-time PCR instrument with a temperature gradient of 25–95 °C at a heating rate of 1 °C/min.

### FcγRIIIa mediated reporter gene assay

SK-BR-3 cells were seeded at 5 × 10^4^ cells/well in 96 well plate and cultured overnight. The next day, the medium was removed, and 100 µL of antibody solution in IMDM at twice the final concentration (200 nM to 2.56 pM, 5-fold dilution series) was added. FcγRIIIa reporter cells (2 × 10^5^ cells) suspended in 100 µL medium were added to each well and incubated for 6 h. After incubation, the 96-well plate was centrifuged at 400 × g for 5 min. Twenty µL of the supernatant was transferred to 96 well black plate, and QUANTI-Luc reagent solution (50 µL) was added. Luminescence was measured immediately after. Cells alone served as the negative control, and 200 nM WT served as the positive control. Luminescence values were normalized to the positive and negative controls (cells only), which were defined as 1 and 0, respectively.

### FcγRIIIa affinity chromatography

FcγRIIIa affinity chromatography was performed using a TSKgel FcR-IIIA-NPR column (TOSOH) with buffer A (50 mM sodium citrate, 150 mM NaCl, pH 6.5) and buffer B (50 mM sodium citrate, 150 mM NaCl, pH 4.5). Antibodies were eluted using the following gradient: 0% B for 2 min, 0–100% B over 18 min, 100% B for 5 min, and 100–0% B over 5 min.

### Flow cytometry

SK-BR-3 cells (1.0×10^5^ cells/well) were resuspended in 100 μl of 2-fold serial dilution series of antibodies in HBSS containing 0.1% sodium azide and 5% FBS and incubated on ice for 45 min. After washing twice, the cells were resuspended in 50 μL of PE-goat anti-human IgG (1/400 dilution, #109–116–170, Jackson ImmunoResearch) in HBBS and incubated on ice for 30 min. The cells were washed once, resuspended in HBBS, and analyzed using a BD LSRFortessa Cell Analyzer (BD Biosciences). Data were analyzed using FlowJo software. MFI values were normalized to the positive (WT 500 nM) and negative (cells only) controls, which were defined as 1 and 0, respectively.

### Surface plasmon resonance (SPR)

SPR measurements were conducted using a Biacore T200 instrument (Cytiva) run in HBS-EP+ at 32 °C. Antibodies were captured on a CM5 chip using a Human Antibody Capture Kit (Cytiva) at a flow rate of 10 µL/min for 30 s to reach approximately 170 RU. HER2-MBP (0.313 nM to 20 nM, 2-fold dilution series) was flowed at 30 µL/min for 180 s, followed by 420 s of dissociation. Antibodies were regenerated by 3 M MgCl_2_ for 30 s at a flow rate of 30 µL/min. The interaction between HER2 and the immobilized antibodies was analyzed using multi-cycle kinetics.

### Statics analysis

Data from the ADCC reporter assay and flow cytometry were fitted to a four-parameter logistic (4PL) model using GraphPad Prism to determine EC50 values, which are presented as mean ± SD. (For the ADCC assay, n = 3 for WT, H-A118C [10, 20, and 40-kDa], and L-V205C [40-kDa]; n = 4 for L-V205C [10 and 20-kDa]. For flow cytometry, n = 3 for all antibody variants.)

## Acknowledgment

This study was partly supported by JSPS (grant number JP24K01270) and AMED (grant numbers JP25ama221234, JP26ama121042, and JP26am0521014) for H.A. Y.T. was supported by JST SPRING (grant number JPMJSP2110). Paperpal was used to assist with grammar and spell checking during the preparation of this manuscript. The authors reviewed all suggested edits and are responsible for the final text.

