## Supporting Information for "Size-dependent effect of site-specific polyethylene glycol modification of trastuzumab on the antibody functions"

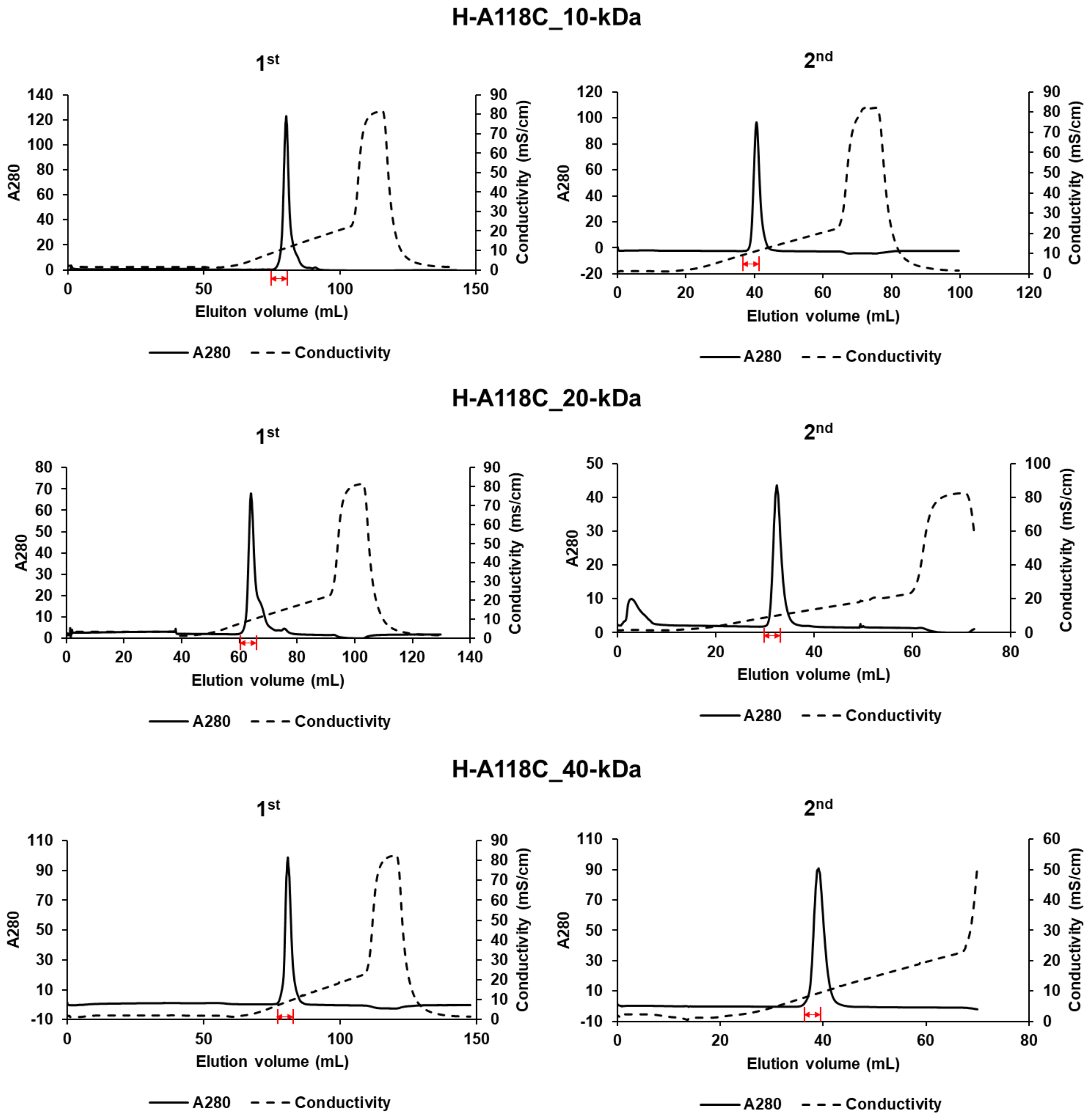


**Figure S1.** (continued to the next page)

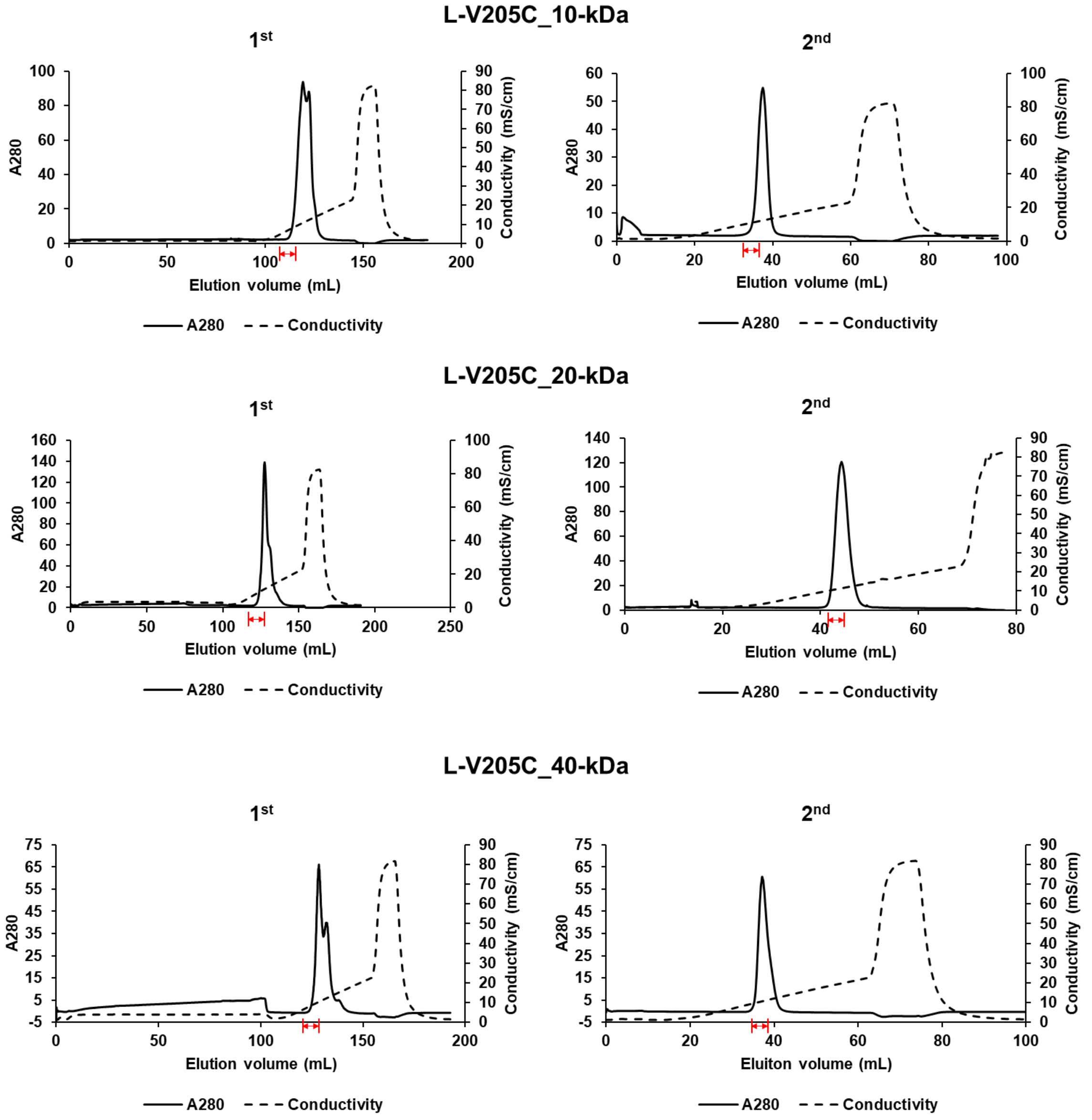


**Figure S1.** Chromatograms of cation exchange chromatography (CIEX) for the purification of bis-PEGylated antibodies. The PEGylated antibody was purified by performing cation exchange CIEX twice. In the 1$^{st}$ CIEX, fractions with red arrows were collected and subjected to 2$^{nd}$ CIEX.

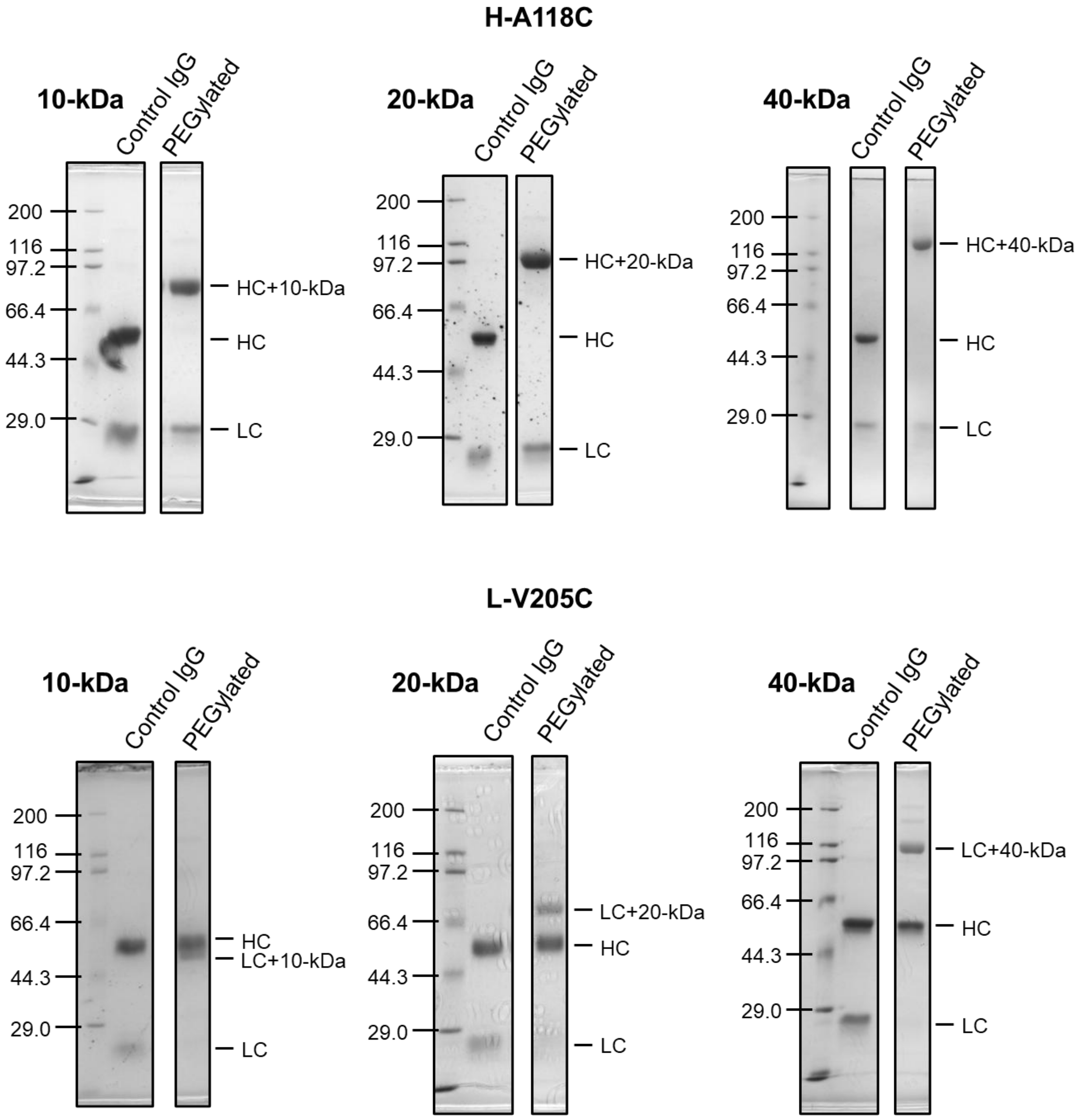


**Figure S2.** SDS-PAGE analysis of purified PEGylated antibodies.

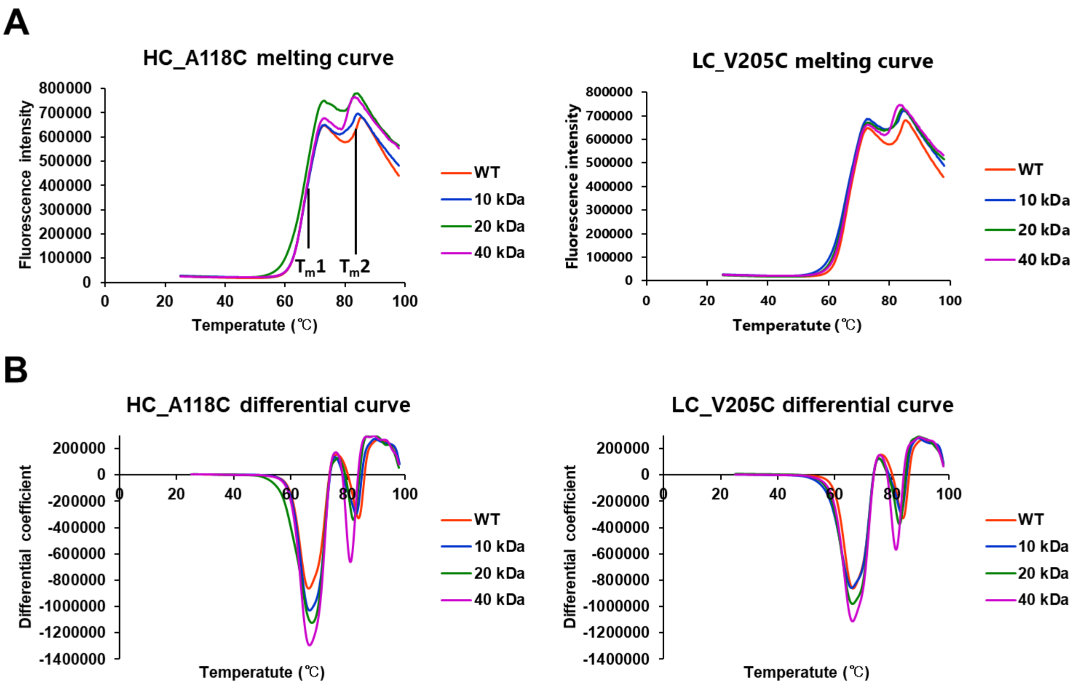


**Figure S3.** Thermal stability of PEGylated antibodies using differential scanning fluorimetry. The melting and differential curves are shown in panels (A) and (B), respectively.

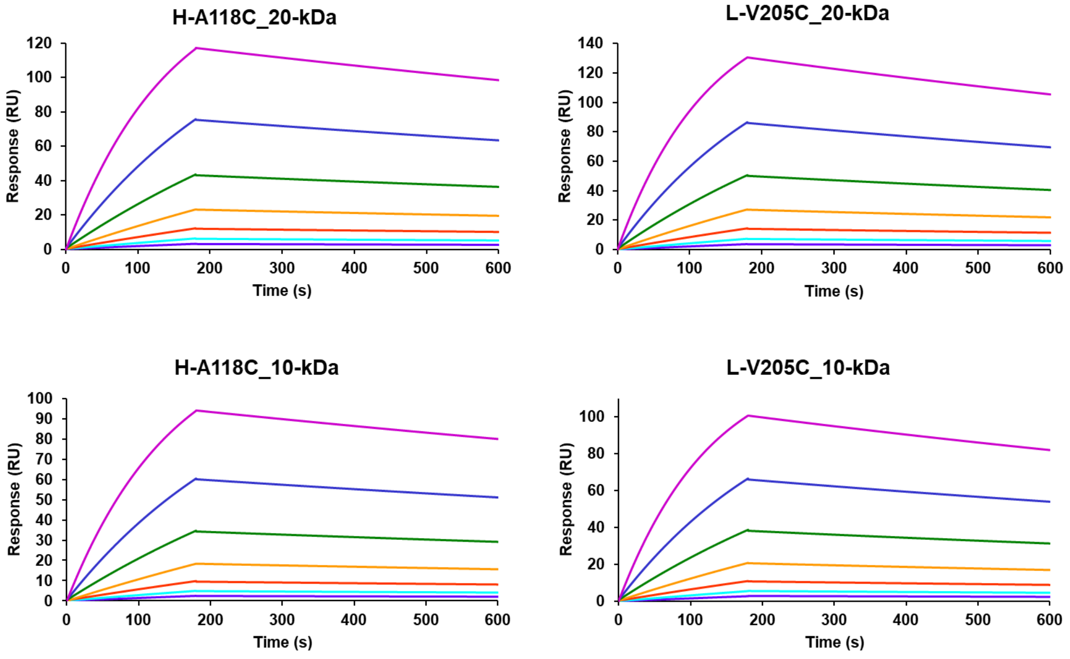


**Figure S4.** Surface plasmon resonance sensorgrams of immobilized PEGylated antibodies (10 or 20-kDa) binding to HER2-MBP as the analyte.

**Table S1.** Melting temperatures in differential scanning fluorimetry[a]

| | | $T_{m1}$ (° C) | $T_{m2}$ (°C) |
|---|---|---|---|
| WT | | 66.2 ± 0.2 | 83.8 ± 0.1 |
| H-A118C | 10-kDa | 66.6 ± 0.2 | 82.7 ± 0.1 |
| | 20-kDa | 67.9 ± 0.4 | 82.3 ± 0.1 |
| | 40-kDa | 66.5 ± 0.1 | 80.9 ± 0.1 |
| L-V205C | 10-kDa | 65.9 ± 0.2 | 83.2 ± 0.2 |
| | 20-kDa | 66.4 ± 0.2 | 82.6 ± 0.1 |
| | 40-kDa | 66.2 ± 0.1 | 81.4 ± 0.2 |

[a] The data are presented as mean ± SD from three independent experiments. The melting temperatures ($T_m$) were determined from the differential curves of the fluorescence intensity profiles.
